# The mechanosensitive channel MscS links lipoteichoic acid synthesis alterations to beta-lactam sensitivity in *Staphylococcus aureus*

**DOI:** 10.64898/2026.05.29.728642

**Authors:** Majid Shah, Amol Kanampalliwar, Taeok Bae

## Abstract

Lipoteichoic acid is an essential cell envelope polymer in *Staphylococcus aureus* and contributes to beta-lactam resistance in methicillin-resistant *S. aureus*. Mutations that alter lipoteichoic acid synthesis sensitize MRSA to beta-lactam antibiotics, but the mechanism connecting envelope polymer defects to antibiotic susceptibility has remained unclear. Here, we used suppressor genetics, genome sequencing, transcriptomics, inducible gene expression, antibiotic susceptibility assays, and measurements of intracellular potassium, cyclic di-AMP, and penicillin-binding protein production to define this connection. We found that lipoteichoic acid synthesis defects increased expression of a ParB-like protein and the mechanosensitive channel MscS. Repression of this locus in RNA polymerase suppressor mutants restored beta-lactam resistance, whereas induced expression of both genes reduced resistance. Increased ParBL-MscS expression was associated with decreased intracellular potassium and reduced cyclic di-AMP, linking altered membrane-envelope physiology to second messenger signaling. Lipoteichoic acid synthesis mutants also showed reduced production of penicillin-binding protein 4, an important determinant of beta-lactam resistance. Modest restoration of penicillin-binding protein 4 improved resistance in the mutant with reduced lipoteichoic acid abundance, whereas the mutant producing elongated lipoteichoic acid showed a distinct response, indicating that different lipoteichoic acid defects impose different envelope stresses. Together, these findings identify a potassium-dependent pathway connecting lipoteichoic acid synthesis, mechanosensitive channel activity, cyclic di-AMP signaling, penicillin-binding protein 4 production, and beta-lactam resistance. This work reveals how cell envelope perturbations are converted into cytoplasmic regulatory responses that control antibiotic susceptibility and suggests that ion homeostasis and cyclic di-AMP signaling may be exploitable pathways for restoring beta-lactam efficacy against MRSA.

**Author Summary:** Antibiotic-resistant *Staphylococcus aureus*, including methicillin-resistant *S. aureus*, is difficult to treat because it can withstand many beta-lactam antibiotics, a widely used class of drugs. Previous work showed that changes in lipoteichoic acid, an important molecule in the bacterial cell envelope, make these bacteria more sensitive to beta-lactams. However, it was not clear how a change at the cell surface affects antibiotic resistance inside the cell. In this study, we found that defects in lipoteichoic acid production activate a pathway involving a predicted membrane channel. This pathway changes the level of potassium inside the bacterial cell and affects a small signaling molecule that helps coordinate cell wall maintenance. These changes also reduce the amount of an enzyme involved in building the cell wall, making the bacteria more vulnerable to beta-lactam antibiotics. Our findings suggest that antibiotic resistance depends not only on the direct targets of antibiotics, but also on how bacteria maintain the balance between their cell envelope, ion levels, and internal signaling. Understanding this connection may help identify new ways to make resistant bacteria more sensitive to existing antibiotics.

## Introduction

*Staphylococcus aureus* is a major human pathogen that causes diseases ranging from skin and soft-tissue infections to pneumonia, endocarditis, sepsis, and other life-threatening invasive infections (1). Treatment is especially challenging when infections are caused by methicillin-resistant *S. aureus* (MRSA), which is resistant to most beta-lactam antibiotics. Because beta-lactams are among the safest and most widely used antibacterial drugs, understanding the mechanisms by which MRSA resists them is important for restoring the activity of existing antibiotics and identifying new targets that weaken bacterial resistance.

The primary determinant of beta-lactam resistance in MRSA is penicillin-binding protein 2a (PBP2a), a cell wall synthesis enzyme encoded by *mecA*. PBP2a has low affinity for most beta-lactams and can continue crosslinking peptidoglycan when other penicillin-binding proteins are inhibited (2–4). However, PBP2a alone does not fully explain MRSA beta-lactam resistance. Full resistance requires coordinated cell envelope synthesis and remodeling, including the activities of additional penicillin-binding proteins, cell wall polymers, and regulatory pathways that maintain envelope homeostasis (5). This indicates that beta-lactam resistance is not only a property of the drug target itself, but also depends on the physiological state of the bacterial cell envelope.

Lipoteichoic acid (LTA) is a major anionic polymer of the Gram-positive cell envelope. In *S. aureus*, LTA is anchored in the cytoplasmic membrane and extends through the cell wall, where it contributes to cell growth, division, ion homeostasis, and envelope integrity (6). LTA is essential for normal growth and viability in *S. aureus*(7, 8). Its synthesis requires three main enzymes: UgtP, which produces the glycolipid anchor diglucosyldiacylglycerol; LtaA, which translocates this glycolipid across the membrane; and LtaS, which polymerizes the phosphoglycerol backbone of LTA (7, 9–11). Mutations affecting these enzymes alter LTA abundance or polymer length and markedly sensitize MRSA to beta-lactam antibiotics(10, 12, 13). Although this phenotype has been reported in multiple studies, the mechanism connecting altered LTA synthesis to reduced beta-lactam resistance has remained unclear.

Several pathways could potentially link LTA defects to beta-lactam susceptibility. PBP2 and PBP4 bind to LTA (14). PBP2 provides glycosyltransferase activity required for peptidoglycan synthesis when PBP2a carries out transpeptidation (15), whereas PBP4 promotes high-level peptidoglycan crosslinking and contributes to full beta-lactam resistance (16, 17). In addition, the second messenger cyclic di-AMP is important for cell wall homeostasis, potassium balance, cell size, wall thickness, and antibiotic resistance (18–20). Increased cyclic di-AMP levels have been associated with enhanced beta-lactam resistance, whereas reduced cyclic di-AMP levels can decrease resistance, in part by increasing production of the cell wall hydrolase LytM(19–25). However, it is not known whether LTA synthesis defects influence these pathways to control beta-lactam resistance.

In this study, we sought to determine how alterations in LTA synthesis reduce beta-lactam resistance in MRSA. Using two distinct LTA synthesis mutants and their suppressor mutants, we identified a previously uncharacterized ion channel pathway linking LTA defects to antibiotic susceptibility.

## Results

### Altered LTA synthesis reduces oxacillin resistance in MRSA

To examine the role of LTA synthesis in beta-lactam resistance, we used two distinct MRSA mutants altered in LTA synthesis: MW2:Pspac-*ltaS* and USA300:*ugtP*. MW2:Pspac-*ltaS* is derived from the MW2 strain in which the native promoter of the LTA synthesis gene *ltaS* is replaced with the IPTG-inducible *Pspac* promoter. In the absence of IPTG, this strain produces markedly reduced levels of LTA (Fig 1B). USA300:*ugtP* is a transposon mutant of *ugtP* and, as previously reported, produces a longer LTA polymer (Fig 1B)(10, 12, 13). Both mutants displayed increased sensitivity to oxacillin (Fig 1C), confirming the essential contribution of intact LTA to MRSA beta-lactam resistance. Growth analysis further revealed that MW2:*Pspac-ltaS* exhibited a mild growth defect in the absence of IPTG, whereas USA300:*ugtP* exhibited a more pronounced growth impairment (Fig 1D).

**Fig 1.**
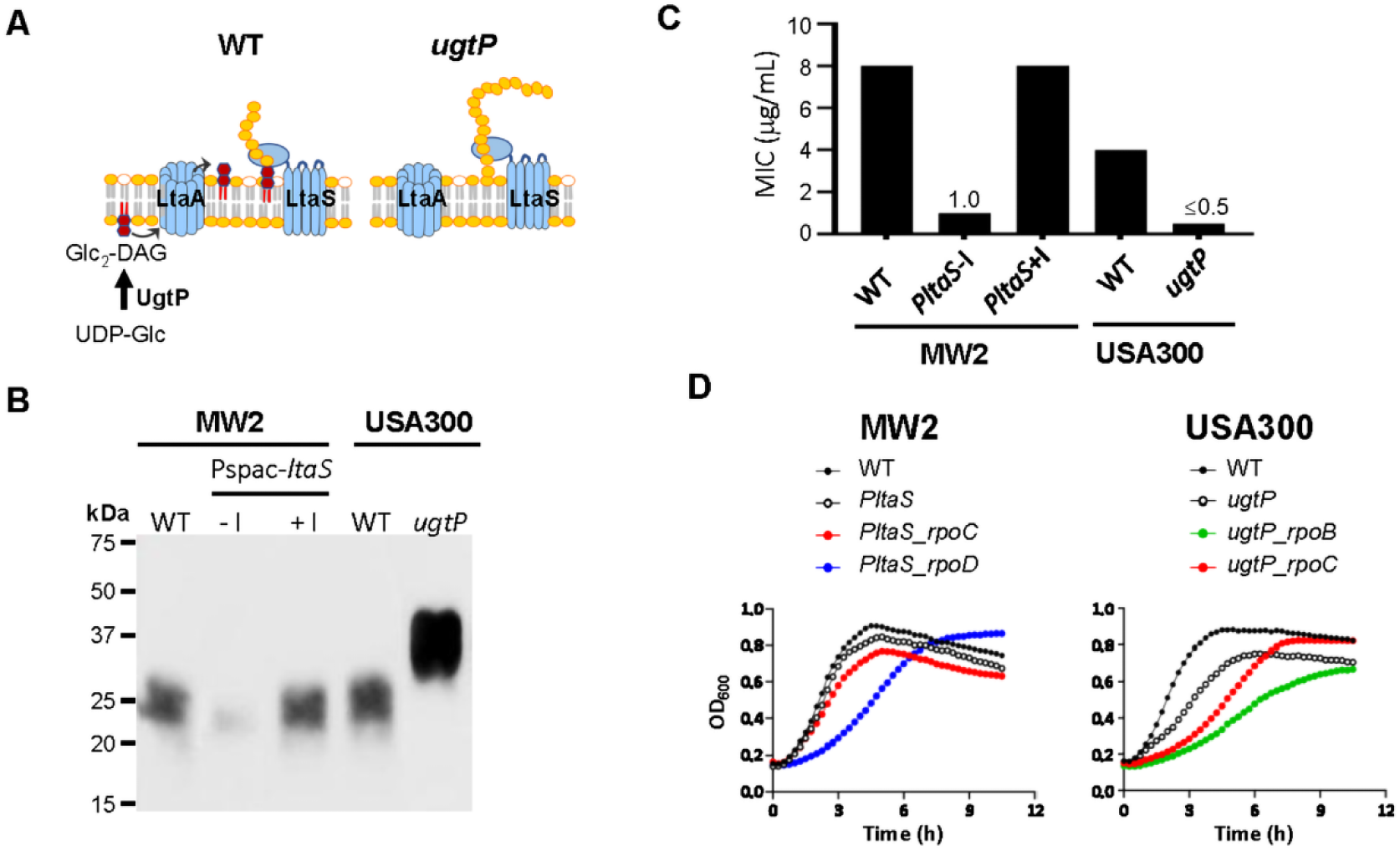
Disruption of LTA synthesis reduces oxacillin resistance and affects growth in MRSA. (A) Schematic representation of the LTA synthesis pathway. Glc₂-DAG, diglucosyldiacylglycerol; UDP-Glc, uridine-5′-diphosphate-glucose. (B) Western blot analysis of LTA production in the indicated LTA synthesis mutants. WT, wild type; Pspac*-ltaS*, MW2:Pspac*-ltaS* grown in the absence of IPTG (–I) or in the presence of 1 mM IPTG (+I). (C) Oxacillin minimum inhibitory concentrations (MICs) of the indicated LTA synthesis mutants. (D) Growth of the LTA synthesis mutants and their suppressor strains in tryptic soy broth (TSB), as measured by OD_600_ over time.

### Identification of suppressor mutations restoring beta-lactam resistance in LTA synthesis mutants

To explore how alterations in LTA synthesis sensitize MRSA to beta-lactams, we performed a suppressor screen. LTA synthesis mutants were plated on tryptic soy agar containing oxacillin at concentrations ranging from 2 to 32 μg/mL. Twelve mutants were selected for each and subjected to genome sequencing. MW2:Pspac-*ltaS* suppressor mutants carried mutations in genes involved in c-di-AMP turnover, including *gdpP* and *pde2*, as well as in RNA polymerase subunit genes *rpoC* and *rpoD* (Table 1 and S1 Table). In contrast, USA300:*ugtP* suppressors harbored mutations in *gdpP, prs*, a gene involved in nucleotide and amino acid (e.g., tryptophan and histidine) biosynthesis (26), and in the RNA polymerase subunit genes *rpoB* and *rpoC* (Table 1 and S1 Table).

**Table 1.** Suppressor mutants isolated in this study.

| No. | Position | Mut. | a.a.<br>change | Gene | Product | Add.<br>Mut. |
| --- | --- | --- | --- | --- | --- | --- |
| **MW2:Pspac-*ltaS*\*** |  |  |  |  |  |  |
| **2.19** | **19325** | **G/A** | **G335D** | ***gdpP*** | **c-di-AMP phosphodiesterase** | **3** |
| **4.10** | **19358** | **C/G** | **P346R** | ***gdpP*** | **c-di-AMP phosphodiesterase** | **7** |
| 2-13 | 1783246 | G/T | S294* | *pde2* | pApA hydrolase | 1 |
| 2-07 | 1783442 | G/A | Q229* | *pde2* | pApA hydrolase | 3 |
| 2-11 | 1783520 | G/A | Q203* | *pde2* | pApA hydrolase | 4 |
| 2-05 | 1784041 | T/C | D29G | *pde2* | pApA hydrolase | 5 |
| 4-05 | 576084 | C/T | R739C | *rpoC* | RNA polymerase, $\beta'$ subunit | 4 |
| **4-15** | **576087** | **G/T** | **G740C** | ***rpoC*** | **RNA polymerase, $\beta'$ subunit** | **0** |
| 4-04 | 576267 | G/T | D800Y | *rpoC* | RNA polymerase, $\beta'$ subunit | 1 |
| 2-14 | 1641080 | C/A | V338F | *rpoD* | Sigma factor 70 | 1 |
| **2-16** | **1641080** | **C/A** | **V338F** | ***rpoD*** | **Sigma factor 70** | **0** |
| 4-03 | 1641121 | C/T | G320D | *rpoD* | Sigma factor 70 | 3 |
| **USA300:*ugtP*** |  |  |  |  |  |  |
| **4-29** | **18680** | **+A** | **F.S.** | ***gdpP*** | **cyclic-di-AMP phosphodiesterase** | **3** |
| **2-16** | **19732** | **G/C** | **S463T** | ***gdpP*** | **cyclic-di-AMP phosphodiesterase** | **4** |
| 32-01 | 535248 | G/C | G35R | *prs* | Phosphoribosylpyrophosphate synthetase | 0 |
| 32-05 | 535411 | C/T | A89V | *prs* | Phosphoribosylpyrophosphate synthetase | 0 |
| 32-08 | 535581 | G/T | D146Y | *prs* | Phosphoribosylpyrophosphate synthetase | 0 |
| 16-05 | 535693 | T/A | V183D | *prs* | Phosphoribosylpyrophosphate synthetase | 0 |
| 16-06 | 535839 | G/A | D232N | *prs* | Phosphoribosylpyrophosphate synthetase | 0 |
| **16-07** | **587109** | **C/A** | **P519H** | ***rpoB*** | **RNA polymerase, $\beta$ subunit** | **0** |
| 8-07 | 587463 | T/C | L637S | *rpoB* | RNA polymerase, $\beta$ subunit | 0 |
| 16-3 | 587468 | G/T | G639C | *rpoB* | RNA polymerase, $\beta$ subunit | 0 |
| 8-04 | 590723 | T/A | D494E | *rpoC* | RNA polymerase, $\beta'$ subunit | 0 |
| **8-10** | **592417** | **A/G** | **Y1059C** | ***rpoC*** | **RNA polymerase, $\beta'$ subunit** | **0** |

Mut., mutation; Add. Mut., additional mutation number; F.S., frame-shift; bold-faced mutants are used in this study.

Suppressor mutations that restored oxacillin resistance were identified in genes involved in cyclic di-AMP metabolism, nucleotide biosynthesis, and RNA polymerase function. Because RNA polymerase mutations occurred in both LTA mutant backgrounds and could reveal transcriptional changes linked to restored resistance, we selected representative RNA polymerase suppressors for further analysis. Four representative suppressor mutants were selected: MW2:*Pspac-ltaS rpoC* G740C (no. 4-15), *rpoD* V338F (no. 2-16), USA300:*ugtP rpoB* P519H (no. 16-07), and *rpoC* Y1059C (no. 8-10) (Table 1 and S1 Table). All altered residues are conserved in *Escherichia coli* RNA polymerase (S1 Fig). In tryptic soy broth, these four suppressor mutants exhibited slower growth than their respective wild-type and parent strains (Fig 1D).

### Transcriptomic profiling reveals three genes consistently downregulated in RNA polymerase suppressor mutants

Because RNA polymerase controls global transcription, we hypothesized that the enhanced beta-lactam resistance of the suppressor mutants results from altered transcription of specific genes common to all four mutants. To identify these, we performed RNA-seq analysis of the RNA polymerase suppressor mutants relative to their respective parent strains, MW2:*Pspac-ltaS* and USA300:*ugtP*. The MW2:Pspac-*ltaS* suppressor mutants displayed widespread transcriptional changes, with over 1,000 differentially expressed genes (*rpoC* mutant: 1,260 genes; *rpoD* mutant: 1,310 genes). In contrast, the USA300:*ugtP* suppressors exhibited fewer changes (*rpoB* mutant: 70; *rpoC* mutant: 202) (Fig 2A). Approximately 80% of differentially expressed genes were shared between the two MW2 suppressors, while only 3–20% overlap was observed between the USA300 mutants (Fig 2A). Despite these global differences, three genes were consistently downregulated across all four suppressor mutants: SAUSA300_0361 (MW0336), encoding a ParB-like putative transcriptional regulator (hereafter *parBL*); SAUSA300_0362 (MW0337), encoding a putative small-conductance mechanosensitive ion channel (*mscS*); and SAUSA300_1243 (MW1233), encoding the DNA repair protein SbcC (Fig 2A and 2B). qRT–PCR confirmed the RNA-seq results: transcript levels of all three genes were elevated in the LTA synthesis mutants and reduced in the suppressor mutants (Fig 2C).

**Fig 2.**
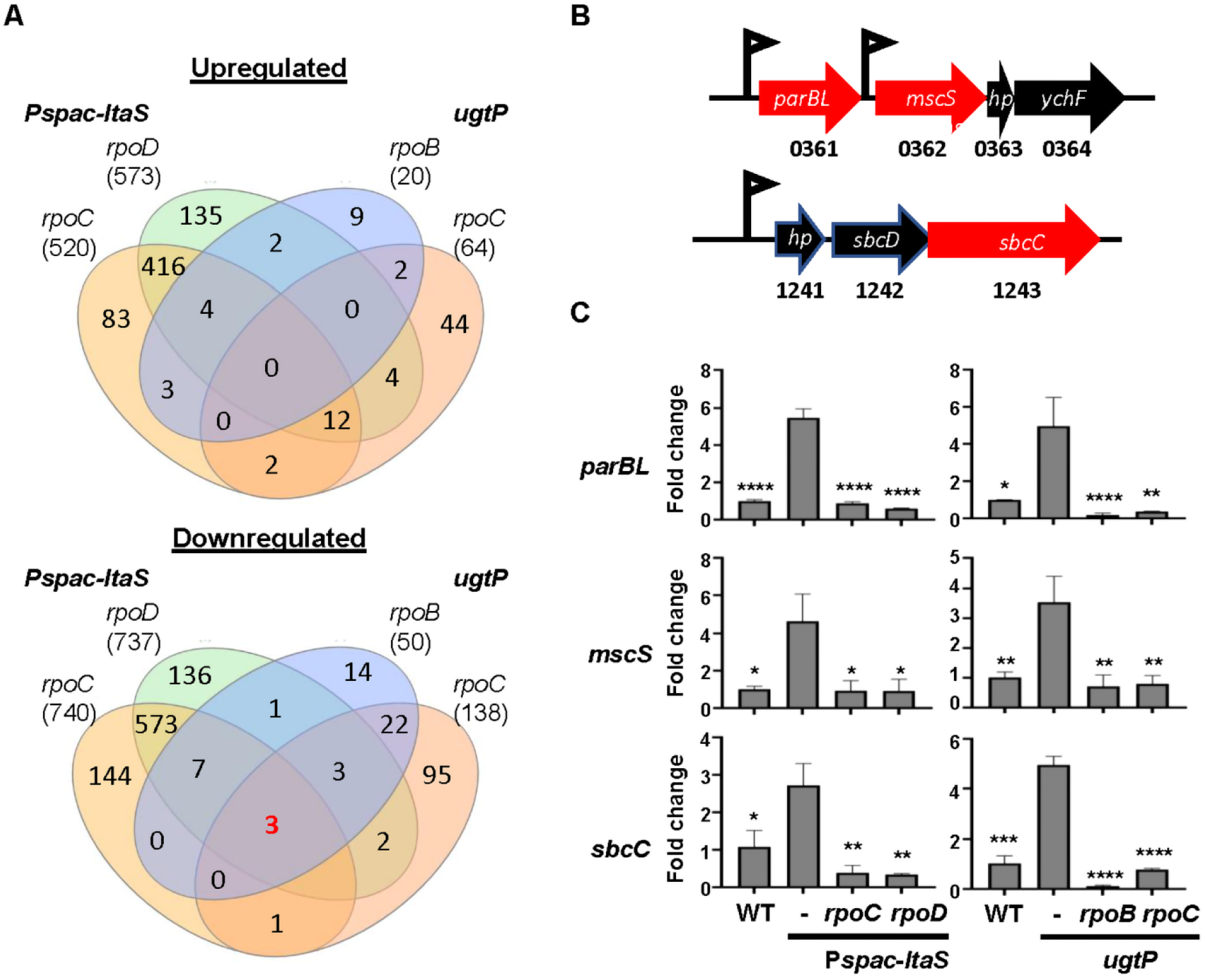
Three genes are consistently downregulated in RNA polymerase suppressor mutants. (A) Comparative RNA-seq analysis of four RNA polymerase suppressor mutants relative to their respective LTA synthesis mutant parent strains. The numbers indicate differentially expressed genes in each comparison. Pspac-*ltaS*, MW2:Pspac*-ltaS*; *ugtP*, USA300:*ugtP*. (B) Genomic organization of the three genes commonly downregulated in all four suppressor mutants. Gene locus numbers correspond to the *S. aureus* USA300 genome, and the three genes are highlighted in red. (C) qRT–PCR validation of the three commonly downregulated genes. “–” indicates strains without suppressor mutations. Statistical significance was determined relative to the corresponding parent LTA synthesis mutants using an unpaired *t*-test. *, *p* < 0.05; **, *p* < 0.005; ***, *p* < 0.0005; ****, *p* < 0.0001.

### Repression of *parBL-mscS* enhances beta-lactam resistance in RNA polymerase suppressor mutants

To identify which of the three downregulated genes contributes to beta-lactam resistance in the suppressor mutants, we sought to inactivate each gene in the LTA synthesis mutant. Both MW2:*Pspac-ltaS* and USA300:*ugtP* carry an erythromycin resistance marker and are not amenable to the transduction of transposon (Tn) mutation (27, 28). Therefore, we generated a clean *ugtP* deletion in USA300 (USA300Δ*ugtP*) and transduced the Tn mutation into the strain. While Tn insertion was successfully introduced into *sbcC* of USA300Δ*ugtP,* we failed to transduce the Tn insertions into *parBL* or *mscS* in the strain. The USA300Δ*ugtP:sbcC* strain showed the same oxacillin MIC as USA300Δ*ugtP* (S2 Fig), indicating that reduced *sbcC* expression does not explain the suppressor phenotype.

Since the transduction approach was not successful for *parBL* and *mscS*, instead, we replaced the native *parBL* promoter with the IPTG-inducible Pspac promoter in both WT and Δ*ugtP* strains of USA300. In the WT background, promoter replacement had no effect on oxacillin susceptibility, regardless of IPTG (Fig 3A and 3B). In USA300Δ*ugtP*, however, Pspac replacement increased the oxacillin resistance in the absence of IPTG, but the addition of IPTG abolished the increase (Fig 3A and 3B; S3 Fig). These results indicate that reduced *parBL-mscS* expression contributes to enhanced beta-lactam resistance in the suppressor mutants. To further examine their roles, *parBL* and *mscS* were cloned individually or together into the multicopy plasmid pOS1-Pitet under an anhydrotetracycline (ATc)-inducible promoter. These plasmid constructs were introduced into the RNA polymerase suppressor mutants, and oxacillin MICs were determined in the presence and absence of ATc. In the presence of ATc, *parBL* and *mscS* alone had a minor effect on oxacillin MIC (S4 Fig). On the other hand, co-expression of both genes markedly reduced oxacillin resistance (4 to 32-fold) (Fig 4A; S5 Fig). Together, these findings demonstrated that elevated expression of *parBL* and the *mscS* operon reduced oxacillin resistance in LTA synthesis mutants, whereas transcriptional repression of these genes enhanced resistance in the suppressor mutants.

**Fig 3.**
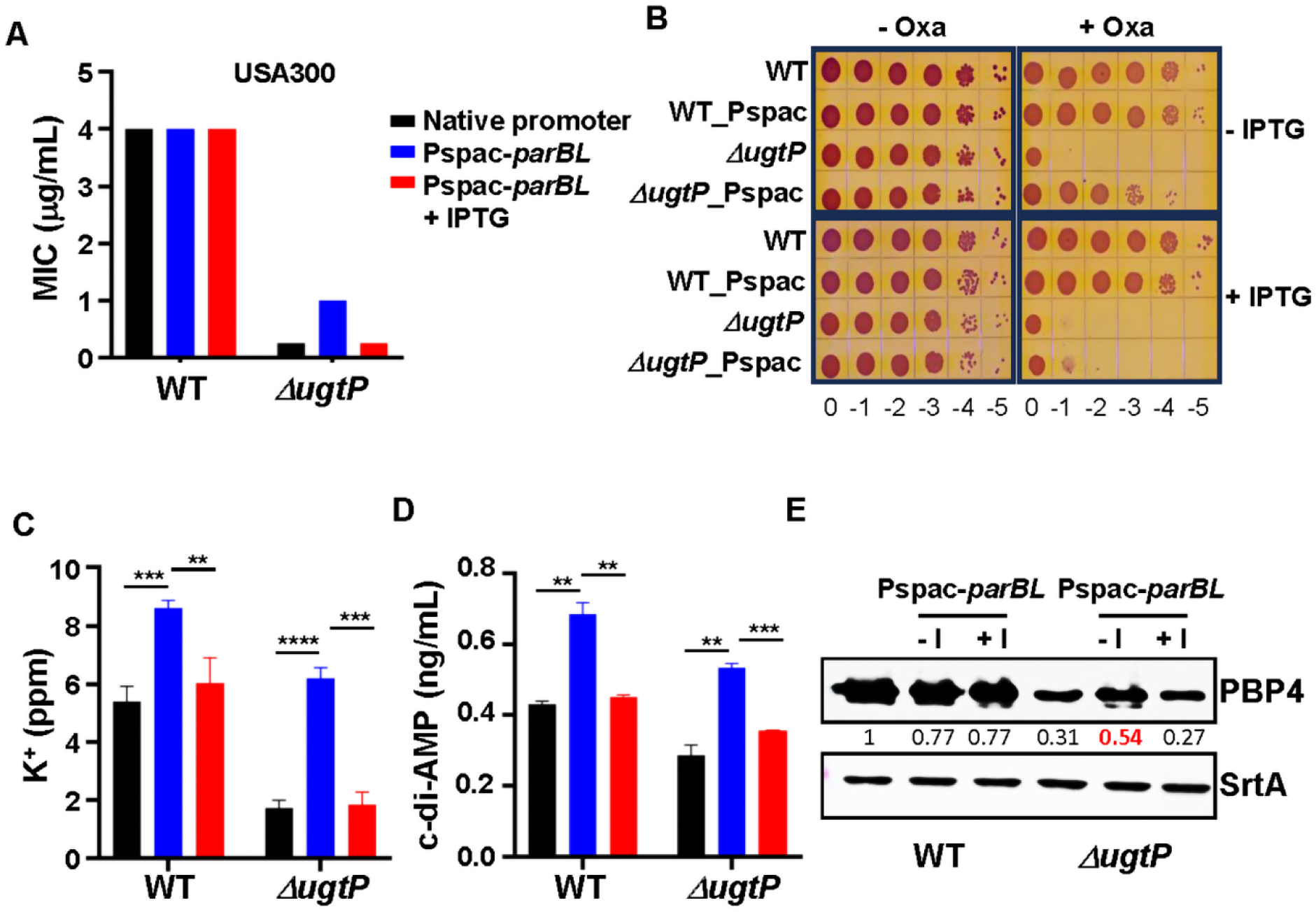
Increased *parBL-mscS* transcription reduces oxacillin resistance in USA300Δ*ugtP*. The native *parBL* promoter was replaced with the IPTG-inducible *Pspac* promoter to control *parBL-mscS* transcription. Transcription was induced by the addition of 1 mM IPTG. The effects of *parBL-mscS* induction were assessed by measuring (A) oxacillin MIC in TSB, (B) oxacillin resistance on agar plates, (C) intracellular potassium concentration, (D) c-di-AMP levels, and (E) PBP4 production. Statistical significance was determined using an unpaired *t-*test. **, *p* < 0.005; ***, *p* < 0.0005; ****, *p* < 0.0001.

**Fig 4.**
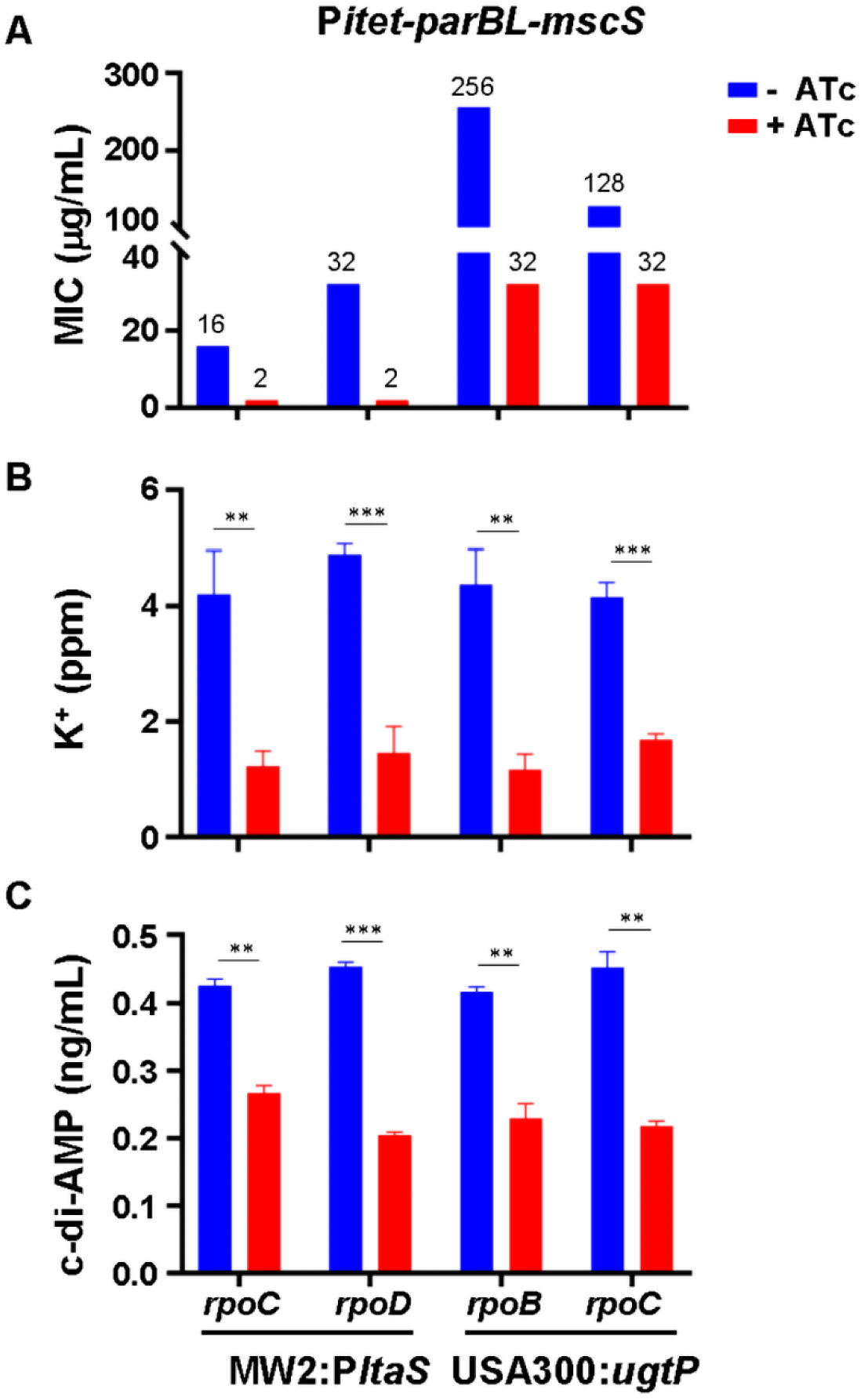
Overexpression of *parBL-mscS* reduces oxacillin resistance, intracellular potassium, and c-di-AMP levels in RNA polymerase suppressor mutants. The *parBL-mscS* locus was overexpressed in the indicated RNA polymerase suppressor mutants using an anhydrotetracycline-inducible promoter. The effects of *parBL-mscS* overexpression were assessed by measuring oxacillin MIC (A), intracellular potassium concentration (B), and intracellular c-di-AMP concentration (C). –ATc, no anhydrotetracycline; +ATc, with anhydrotetracycline (100 ng/mL). Statistical significance was determined using an unpaired *t*-test. **, *p* < 0.005; ***, *p* < 0.0005.

### ParBL/MscS modulates intracellular potassium levels in *S. aureus*

The *mscS* gene is predicted to encode a mechanosensitive ion channel. In *Escherichia coli*, MscS mediates the efflux of chloride and potassium ions from the cytoplasm (29). To determine whether *parBL* and *mscS* have similar functions in *S. aureus*, we measured intracellular potassium levels in the relevant mutants. Intracellular potassium was markedly reduced in USA300Δ*ugtP* compared with the WT strain (Fig 3C). In Pspac*-parBL* fusion strains, potassium levels were elevated in the absence of IPTG and decreased upon IPTG induction (Fig 3C). Consistent results were obtained in the *parBL-mscS* overexpression experiments with the RNA polymerase suppressor mutants: ATc-induced expression of *parBL* and *mscS* significantly lowered intracellular potassium levels (Fig 4B). These results indicated that ParBL and MscS modulated intracellular potassium concentration, likely through MscS-mediated ion efflux, and that beta-lactam resistance in the LTA synthesis mutants is influenced by this potassium-dependent regulatory pathway.

### Intracellular potassium modulates c-di-AMP signaling in *S. aureus*

Intracellular potassium and the second messenger c-di-AMP are tightly linked in several Gram-positive bacteria, including *Bacillus subtilis* and *Streptococcus pneumoniae*, where elevated potassium levels promote c-di-AMP synthesis (30, 31). To determine whether this relationship also exists in *S. aureus*, we analyzed the correlation between intracellular potassium and c-di-AMP concentrations. In the suppressor mutants, overexpression of *parBL-mscS* significantly reduced c-di-AMP levels, showing a positive correlation between intracellular potassium and c-di-AMP (Fig 4B and 4C). We next examined this relationship in the LTA synthesis mutants and their suppressors. Promoter–*lacZ* fusion assays revealed that *parBL* and *mscS* promoter activities were elevated in the LTA synthesis mutants and reduced in their suppressors, indicating transcriptional regulation (Fig 5A and 5B). Intracellular potassium levels were inversely correlated with *parBL* and *mscS* promoter activities (Fig 5C), while c-di-AMP levels again positively correlated with intracellular potassium (Fig 5D). Because c-di-AMP in *S. aureus* is synthesized by the diadenylate cyclase DacA (19), we next tested whether potassium-dependent modulation of c-di-AMP occurs at the transcriptional level. A *dacA* promoter–*lacZ* fusion showed that beta-galactosidase activity correlated with potassium levels across the mutant and suppressor strains (Fig 5E), suggesting that intracellular potassium promotes c-di-AMP production by upregulating *dacA* transcription. These data showed a positive association among intracellular potassium levels, *dacA* promoter activity, and c-di-AMP abundance in the LTA synthesis mutants and suppressor strains.

**Fig 5.**
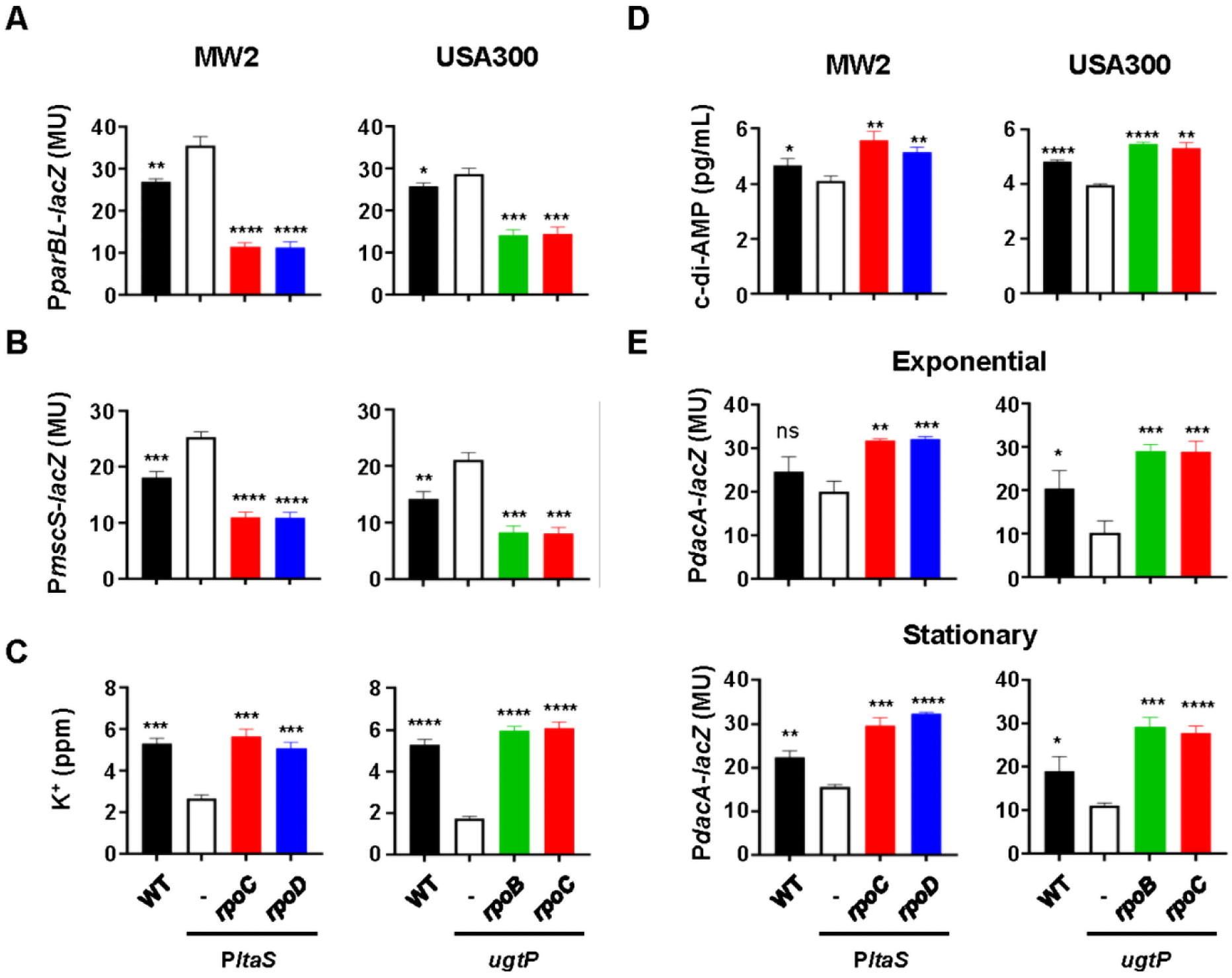
Increased parBL and mscS promoter activities are associated with reduced intracellular potassium, reduced c-di-AMP levels, and decreased *dacA* promoter activity. (A) *parBL* promoter activity measured using a promoter–*lacZ* fusion assay. (B) *mscS* promoter activity measured using a promoter–*lacZ* fusion assay. (C) Intracellular potassium concentration. (D) Intracellular c-di-AMP concentration. (E) *dacA* promoter activity measured using a promoter–*lacZ* fusion assay during exponential and stationary growth phases. Statistical significance was determined relative to the corresponding LTA synthesis mutant parent strain using an unpaired *t-*test. *, *p* < 0.05; **, *p* < 0.005; ***, *p* < 0.0005; ****, *p* < 0.0001.

### LTA synthesis defects reduce PBP4 production and alter PBP4-dependent oxacillin resistance

Penicillin-binding proteins (PBPs) are the molecular targets of beta-lactam antibiotics. To determine whether the altered oxacillin susceptibility of the LTA synthesis mutants and their suppressors is associated with changes in PBP expression, we analyzed cellular PBP levels by Western blotting. Intriguingly, only PBP4 expression was markedly reduced in the LTA synthesis mutants and fully or partially restored in the suppressor mutants (Fig 6A).

**Fig 6.**
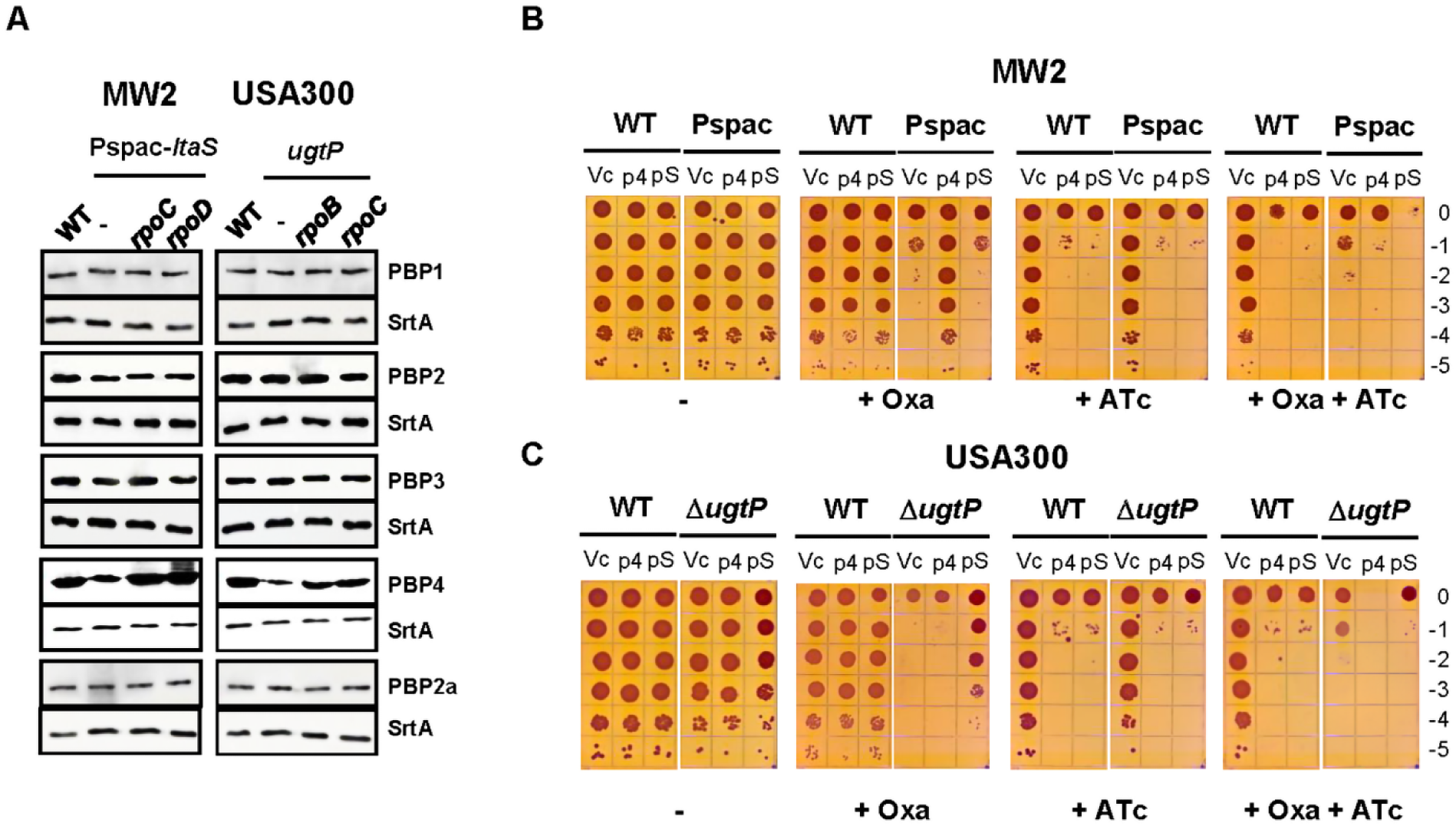
LTA synthesis mutations reduce PBP4 expression and alter the effect of PBP4 on oxacillin resistance. (A) Western blot analysis of penicillin-binding proteins (PBPs) in the indicated strains. Sortase A (SrtA) was used as a loading control. (B) Effect of PBP4 expression on oxacillin resistance in MW2 wild-type (WT) and MW2:Pspac-*ltaS* (Pspac) strains, assessed by spot dilution assay. (C) Effect of PBP4 expression on oxacillin resistance in USA300 WT and Δ*ugtP* strains, assessed by spot dilution assay. The numbers indicate serial 10-fold dilutions from 10⁰ to 10⁻⁵. Vc, pOS1 empty vector; p4, pOS1-Pitet-*pbp4*; pS, pOS1-Pitet-*pbp4 S75A*; –, no antibiotic; + Oxa, oxacillin added; + ATc, anhydrotetracycline added.

To examine whether the decreased PBP4 levels are responsible for the reduced beta-lactam resistance of the LTA synthesis mutants, we generated a PBP4 expression plasmid pOS1-Pitet-*pbp4*, in which the *pbp4* gene is transcribed by the ATc-inducible promoter, and transformed WT and the LTA synthesis mutants with the plasmid. In the LTA synthesis mutants, the basal expression of PBP4 from the plasmid increased PBP4 levels to 51% of the WT level (S6 Fig). On the other hand, when ATc was added, PBP4 levels were increased to 166% (MW2:Pspac-*ltaS*) or 108% (USA300Δ*ugtP*) of WT levels (S6 Fig). Intriguingly, when PBP4 was overexpressed, bacterial growth was inhibited (S7 Fig), indicating that PBP4 overexpression is toxic to the cells.

Next, we examined whether increased PBP4 expression could restore oxacillin resistance using the spot dilution assay. In the absence of ATc, the PBP4 expression plasmid restored oxacillin resistance in MW2:Pspac*-ltaS* (+ Oxa, p4 in Fig 6B), suggesting that the decreased PBP4 level in MW2:Pspac*-ltaS* is responsible for the reduced oxacillin resistance. The replacement of the active-site serine with alanine in PBP4 abolished the restoration (+ Oxa, pS in Fig 6B), suggesting that the increased PBP4 activity, not the protein itself, is responsible for the restored oxacillin resistance. However, when PBP4 expression was induced by ATc, bacterial growth was inhibited in both WT and Pspac-*ltaS* strains, regardless of the presence of oxacillin (+ ATc and + Oxa + ATc in Fig 6B), further confirming that PBP4 overexpression is toxic to the strains. Notably, growth inhibition was also observed with the active-site mutant of PBP4 (pS, + ATc and + Oxa + ATc in Fig 6B), suggesting that the protein itself, not the enzymatic activity, is responsible for the growth inhibition.

In the USA300Δ*ugtP* strain, however, we observed different results. In the absence of ATc, the basal expression of PBP4 failed to restore oxacillin resistance (+ Oxa, p4 in Fig 6C). Unexpectedly, the active-site mutant of PBP4 partially restored oxacillin resistance (+ Oxa, pS in Fig 6C). As with MW2 strains, the ATc-mediated induction of PBP4 expression inhibited the growth of the strains, regardless of the active site mutation in PBP4. The distinct effects of basal PBP4 expression on oxacillin resistance in MW2:Pspac-*ltaS* and USA300Δ*ugtP* suggest that reduced LTA abundance and production of elongated LTA polymers elicit different types of cell envelope stress.

### ParBL/MscS negatively regulates PBP4 expression at the post-transcriptional level through potassium-dependent signaling

Both PBP4 and c-di-AMP are critical for beta-lactam resistance in *S. aureus*. Because *parBL-mscS* expression negatively affects c-di-AMP levels, likely through modulation of intracellular potassium, we next examined whether this regulatory module also influences PBP4 expression. We analyzed PBP4 levels in USA300Δ*ugtP* carrying a Pspac*-parBL* promoter replacement. In the wild-type strain, IPTG induction did not affect PBP4 expression (Fig 3E). In contrast, in the Δ*ugtP* background, PBP4 levels were elevated in the absence of IPTG but decreased upon IPTG addition (Fig 3E). However, *pbp4* transcription was not significantly affected by IPTG (S8 Fig). These results indicate that ParBL/MscS negatively regulates both c-di-AMP and PBP4 expression through intracellular potassium in the context of LTA synthesis defects.

### *gdpP* suppressor mutations increase c-di-AMP and differentially affect potassium, PBP4, and LytM

Reduced expression of ParBL/MscS increased intracellular potassium levels and was accompanied by elevated c-di-AMP and PBP4 levels (Fig 3D and 3E). In *S. aureus*, however, c-di-AMP is a central regulator of potassium homeostasis, primarily through inhibition of potassium uptake (32–34). These observations suggest reciprocal regulation between potassium and c-di-AMP. We therefore asked whether increased c-di-AMP levels affect intracellular potassium concentrations and PBP4 expression. To address this question, we analyzed the *gdpP* suppressor mutants (Table 1 and S1 Table). Because a recent study reported that chemical activation of GdpP increases expression of the cell wall hydrolase LytM and thereby reduces beta-lactam resistance in MRSA (25), we also examined whether c-di-AMP affects LytM expression in the LTA mutant backgrounds.

Using a riboswitch reporter plasmid to measure c-di-AMP levels (35), we found that all *gdpP* suppressor mutants had elevated c-di-AMP levels (Fig 7A). These mutants also exhibited higher intracellular potassium levels than their respective parental LTA synthesis mutants (Fig 7B), although the extent of restoration varied by strain background. In the MW2:*P*spac-*ltaS* background, potassium levels remained below those of the WT strain, whereas in the USA300:*ugtP* background, potassium levels were restored to approximately WT levels.

**Fig 7.**
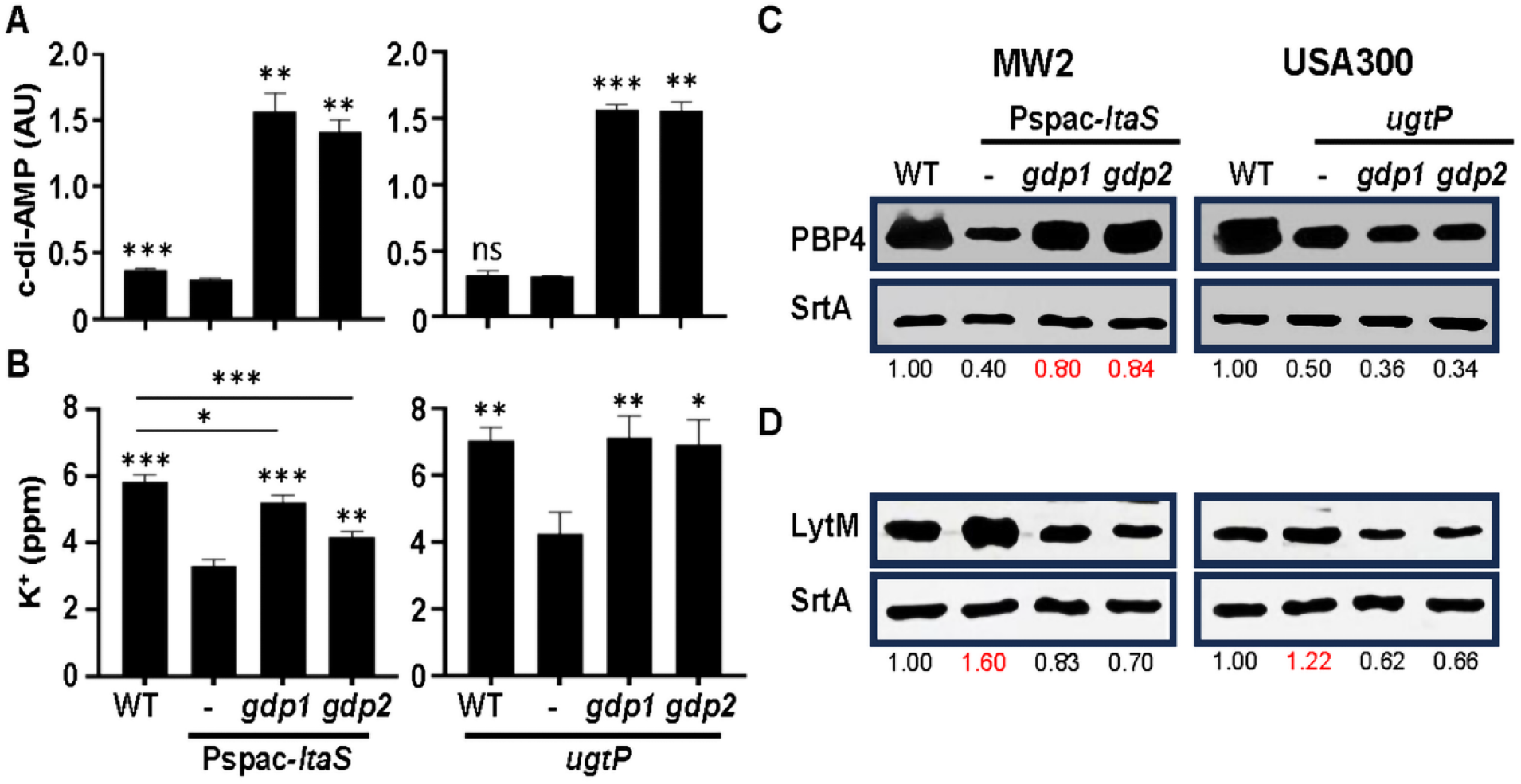
Elevated c-di-AMP levels affect intracellular potassium levels, PBP4 expression, and LytM expression in an LTA mutation- and/or strain-dependent manner. WT strains, LTA synthesis mutants, and their corresponding *gdpP* suppressor mutants were grown in TSB to exponential phase. (A) Relative c-di-AMP levels measured using the *kimA* riboswitch reporter plasmid. (B) Intracellular potassium concentrations. (C) PBP4 production analyzed by Western blotting with an anti-PBP4 antibody. (D) LytM production analyzed by Western blotting with an anti-LytM antibody. Statistical significance was determined relative to the corresponding LTA synthesis mutant strain using an unpaired *t-*test. *, *p* < 0.05; **, *p* < 0.005; ***, *p* < 0.0005.

We next examined PBP4 production. In *gdpP* suppressor mutants derived from MW2:*P*spac-*ltaS*, the reduced PBP4 level was restored to near-WT levels (Fig 7C). In contrast, PBP4 levels remained reduced in *gdpP* suppressor mutants derived from USA300:*ugtP* (Fig 7C). Thus, the effect of c-di-AMP on PBP4 production appears to depend on LTA status, strain background, or both.

LytM expression was broadly consistent with the findings of the previous study (25). In MW2:*P*spac-*ltaS*, LytM expression was higher than in the WT strain, but this increase was abolished by the *gdpP* suppressor mutations (Fig 7D). In USA300:*ugtP*, LytM expression was modestly increased relative to the WT strain, and the *gdpP* suppressor mutations reduced its expression (Fig 7D). Together, these results indicate that elevated c-di-AMP is associated with increased intracellular potassium levels in both strain backgrounds, but its effects on PBP4 and LytM differ by genetic context. In MW2:*P*spac-*ltaS*, elevated c-di-AMP correlated with restoration of PBP4 production and reduced LytM expression, whereas in USA300:*ugtP*, it correlated with reduced LytM expression but did not restore PBP4 production.

It should be noted that, unlike the RNA polymerase suppressor mutants, the *gdpP* suppressor mutants contain three to seven additional mutations (Table 1 and S1 Table). Therefore, we cannot exclude the possibility that these additional mutations contribute to the observed effects on intracellular potassium levels and PBP4 or LytM expression.

### LytM plays a minor role in the reduced oxacillin resistance of LTA synthesis mutants

We next asked whether increased LytM expression contributes to the reduced oxacillin resistance observed in LTA synthesis mutants. To test this, we deleted *lytM* in the WT strain and in the LTA synthesis mutants and then measured oxacillin resistance. In TSB, deletion of *lytM* did not alter oxacillin resistance of the WT strain and the LTA synthesis mutants (S9A Fig). In TSA, *lytM* deletion resulted in only a modest increase in oxacillin resistance (S9B Fig). Together, these results suggest that increased LytM expression makes, at most, a minor contribution to the reduced oxacillin resistance of LTA synthesis mutants.

## Discussion

LTA is essential for cell envelope integrity and contributes to beta-lactam resistance in *S. aureus*. Previous studies have shown that defects in LTA synthesis sensitize MRSA to beta-lactam antibiotics, but the mechanism underlying this phenotype remains poorly understood. In this study, we identified a potassium-dependent pathway that links altered LTA synthesis to beta-lactam susceptibility. Our data support a model in which LTA synthesis defects increase expression of the ParBL/MscS module, reduce intracellular potassium levels, decrease cyclic di-AMP production, lower PBP4 abundance, and increase beta-lactam sensitivity. Thus, LTA appears to influence beta-lactam resistance not only by supporting cell envelope architecture, but also by helping maintain ion homeostasis and downstream regulatory pathways required for cell wall stress tolerance.

A central finding of this study is that ParBL/MscS connects LTA defects to altered potassium homeostasis. Expression of *parBL* and *mscS* was increased in LTA synthesis mutants and reduced in RNA polymerase suppressor mutants that restored oxacillin resistance (Fig 2). Repression of this locus increased oxacillin resistance in the *ugtP* mutant (Fig 3), whereas induced expression of both *parBL* and *mscS* reduced resistance in suppressor strains (Fig 4). These effects correlated inversely with intracellular potassium levels, suggesting that increased ParBL/MscS activity lowered cytoplasmic potassium, likely through direct ion efflux by MscS. Because MscS is predicted to encode a mechanosensitive channel, these findings suggest that LTA-associated envelope stress may be transmitted across the membrane through changes in ion flux.

Our results also extend the current model of cyclic di-AMP signaling in *S. aureus*. Cyclic di-AMP is known to regulate potassium homeostasis and beta-lactam resistance, but our data suggest that potassium levels may also influence cyclic di-AMP production. In LTA synthesis mutants and suppressor strains, intracellular potassium positively correlated with cyclic di-AMP abundance and with *dacA* promoter activity (Fig 3-5). These findings support a model of reciprocal regulation in which cyclic di-AMP controls potassium transport, while potassium availability feeds back on cyclic di-AMP synthesis. This relationship may allow *S. aureus* to coordinate cell wall synthesis with changes in membrane and ion homeostasis during envelope stress.

PBP4 emerged as an important downstream factor in this pathway. LTA synthesis mutants showed reduced PBP4 production, and this reduction was reversed in several suppressor backgrounds (Fig 6 and Fig 7). In the strain with reduced LTA abundance, modest PBP4 expression restored oxacillin resistance (Fig 6B), indicating that decreased PBP4 contributed directly to beta-lactam hypersensitivity. However, PBP4 expression did not restore resistance in the *ugtP* mutant producing elongated LTA, and an active-site mutant of PBP4 partially improved resistance in that background. These observations suggest that different LTA defects impose distinct envelope stresses. Reduced LTA abundance may compromise beta-lactam resistance partly by lowering PBP4-dependent cell wall crosslinking, whereas elongated LTA may perturb envelope organization in a way that cannot be corrected simply by restoring PBP4 catalytic activity. The activity-independent effect of PBP4 also raises the possibility that PBP4 contributes to envelope organization or cell wall synthesis complex stability, in addition to its enzymatic role.

The relationship between cyclic di-AMP, PBP4, and LytM was more context-dependent. Mutations in *gdpP*, which are expected to elevate cyclic di-AMP levels, increased potassium levels and reduced LytM expression in both LTA mutant backgrounds. In the reduced-LTA mutant, these changes were accompanied by restoration of PBP4 production, whereas in the elongated-LTA mutant, PBP4 levels remained low. These findings suggest that cyclic di-AMP-dependent regulation of cell wall homeostasis differs according to the type of LTA defect and possibly the strain background. Because the *gdpP* suppressor strains contained additional mutations, further work will be required to define which phenotypes are directly caused by increased cyclic di-AMP.

This study changes the way LTA-dependent beta-lactam resistance can be viewed. The prevailing model emphasizes LTA as a structural cell envelope polymer needed for normal growth, division, and envelope integrity. Our findings support an expanded model in which LTA also helps maintain the physiological balance between membrane stress, potassium homeostasis, cyclic di-AMP signaling, and peptidoglycan synthesis. In this model, beta-lactam resistance depends not only on PBP2a and peptidoglycan crosslinking, but also on the ability of the cell to preserve ionic and regulatory homeostasis during cell wall stress.

Several important questions remain. First, the mechanism by which LTA defects increase *parBL-mscS* expression is unknown. Because LTA is located outside the cytoplasmic membrane, altered LTA synthesis probably affects *parBL-mscS* indirectly through membrane or envelope stress. Identifying the regulatory factors that control this locus will be important. Second, the biochemical function of ParBL remains unclear. Although annotated as a ParB-like protein, ParBL appears structurally distinct from canonical chromosome-partitioning ParB proteins (S10 Fig), suggesting that it may have a different role. Future experiments should determine whether ParBL regulates *mscS* expression, modulates MscS activity, binds DNA, or interacts with membrane-associated proteins. Third, direct transport assays will be needed to determine whether MscS exports potassium in *S. aureus* and whether channel activity is required for its effect on beta-lactam resistance. Finally, the mechanism by which potassium and cyclic di-AMP influence PBP4 abundance should be defined, including whether regulation occurs through protein stability, translation, localization, or indirect effects on envelope stress.

In summary, our study identifies a regulatory pathway linking LTA synthesis to beta-lactam resistance in MRSA. Defects in LTA synthesis increased ParBL/MscS activity, reduced intracellular potassium, decreased cyclic di-AMP signaling, altered PBP4 and LytM expression, and sensitized cells to beta-lactam antibiotics. These findings reveal how changes in a cell envelope polymer can be converted into cytoplasmic signals that control antibiotic susceptibility. More broadly, they suggest that disrupting the coordination between envelope integrity, ion homeostasis, and second-messenger signaling may provide new strategies to potentiate beta-lactam activity against MRSA.

## Materials and Methods

### Bacterial strains and culture conditions

All bacterial strains and plasmids used in this study are listed in Table 2. *Escherichia coli* strains were cultured in lysogeny broth (LB), and *S. aureus* strains were cultured in tryptic soy broth (TSB) with shaking at 200 rpm. When required, antibiotics and inducers were added at the following final concentrations: ampicillin, 100 μg/mL; erythromycin, 10 μg/mL; chloramphenicol, 10 μg/mL; isopropyl β-D-1-thiogalactopyranoside (IPTG), 1 mM; and anhydrotetracycline (ATc), 100 ng/mL.

**Table 2.** Bacterial strains and plasmids used in this study.

| Strain/plasmid | Characteristics | Source |
| --- | --- | --- |
| *E. coli* DH5α | Restriction-deficient, plasmid-free | NEB |
| *E. coli* BL21(DE3) | F<sup>-</sup> *ompT hsd<sub>B</sub>*(r<sub>B</sub><sup>-</sup> m<sub>B</sub><sup>-</sup>) *gal dcm* (λDE3), lon<sup>-</sup> | NEB |
| *S. aureus* |  |  |
| RN4220 | Restriction-deficient, prophage-cured | (36) |
| MW2 | Clinical isolate, USA400 | NARSA* |
| MW2:Pspac-*ltaS* | The *ltaS* promoter in MW2 was replaced with the Pspac promoter | This study |
| MW2:Pspac-*ltaS_rpoC* | MW2:Pspac-*ltaS* with a suppressor mutation in *rpoC* G740C | This study |
| MW2:Pspac-*ltaS_rpoD* | MW2:Pspac-*ltaS* with a suppressor mutation in *rpoD* V338F | This study |
| USA300-0114 | USA300-1114, a clinical MRSA strain | NARSA |
| USA300-P23 | USA300-0114, where the plasmids 2 and 3 were eliminated. | (37) |
| USA300:*ugtP* | USA300-P23 with transposon insertion in *ugtP* | (13) |
| USA300:*ugtP_rpoB* | USA300:*ugtP* with the suppressor mutation *rpoB* P519H | This study |
| USA300:*ugtP_rpoC* | USA300:*ugtP* with the suppressor mutation *rpoC* Y1059C | This study |
| USA300Δ*ugtP* | USA300-P23 with *ugtP* deletion | This study |
| USA300Δ*ugtP*_Pspac-*parBL* | The promoter for *parBL* was replaced with the Pspac promoter using pMutin-HA | This study |
| Plasmids |  |  |
| pMutin-HA | A suicide plasmid | Bacillus Genetic Stock Center, Columbus, OH |
| pOS1 | A *E. coli*/*S. aureus* shuttle vector, Amp<sup>r</sup>, Cm<sup>r</sup> | (38) |
| pOS1-Pitet-*parBL* | The *parBL* gene was cloned under Pitet of pOS1-Pitet | This study |
| pOS1-Pitet-*mscS* | The *mscS* gene was cloned under the Pitet of pOS1-Pitet | This study |
| pOS1-Pitet-*parBL-mscS* | The *parBL/mscS* genes were cloned under the Pitet of pOS1-Pitet | This study |
| pET28a-*lytM* | *lytM* was cloned into pET28a | This study |

\* Network on Antimicrobial Resistance in *Staphylococcus aureus*

### Replacement of the *parBL* promoter with an IPTG-inducible promoter

To generate an *S. aureus* strain with IPTG-inducible *parBL* expression, the first 512 bp of *parBL*, including its ribosome-binding site, were PCR-amplified from USA300 chromosomal DNA using Phusion DNA polymerase (NEB) and primers P6251/P6252 (Table 3). The PCR product was purified using a QIAquick PCR Purification Kit (Qiagen). The pMutin-HA vector was digested with *Hind*III and *Kpn*I, and the amplified *parBL* fragment was inserted downstream of the IPTG-inducible Pspac promoter by Gibson assembly (39). The resulting plasmid, pMutin-HA-*parBL*, was first electroporated into *S. aureus* RN4220 and subsequently transduced into USA300 wild-type (WT) and Δ*ugtP* strains using phage φ85.

**Table 3.**
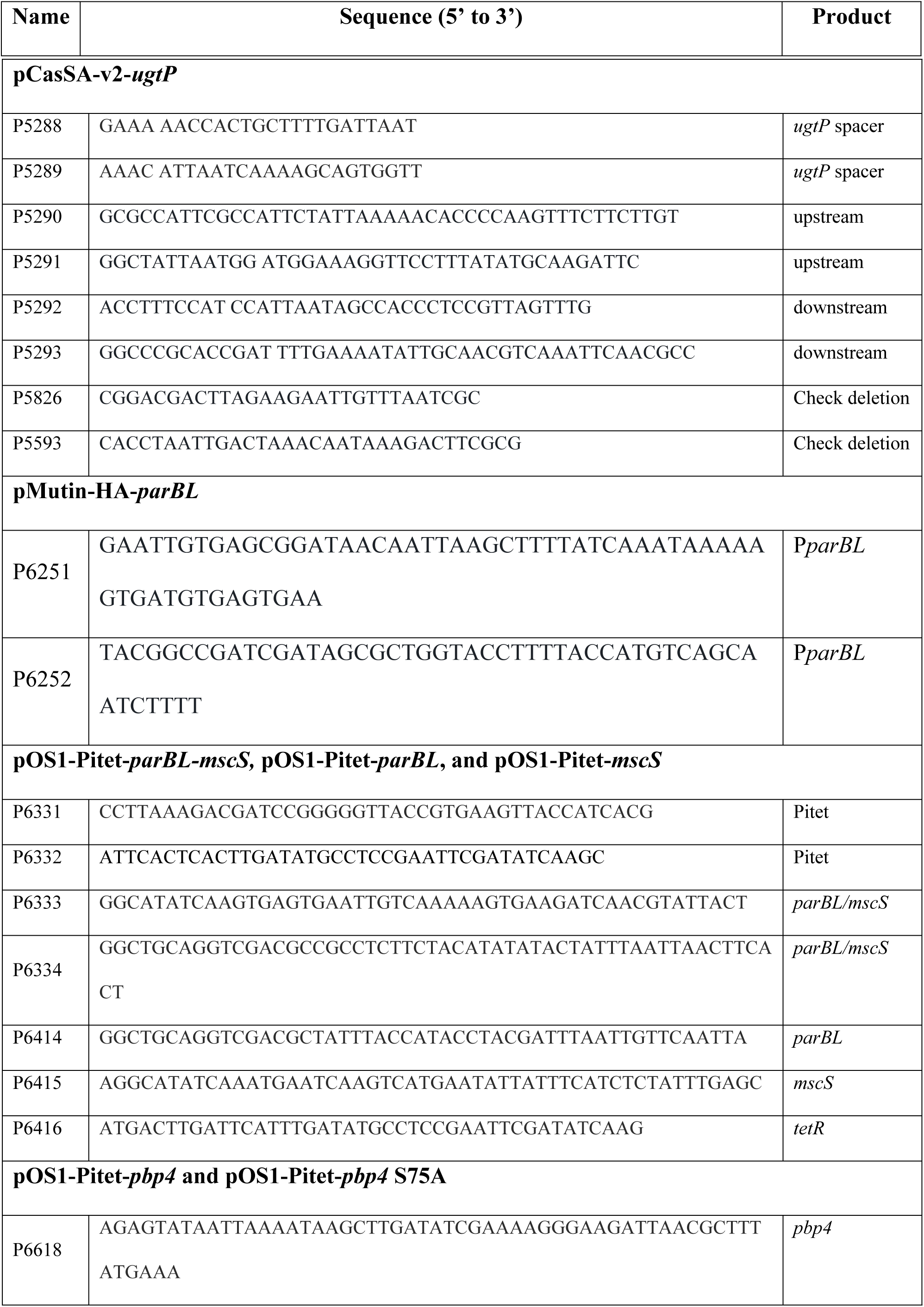
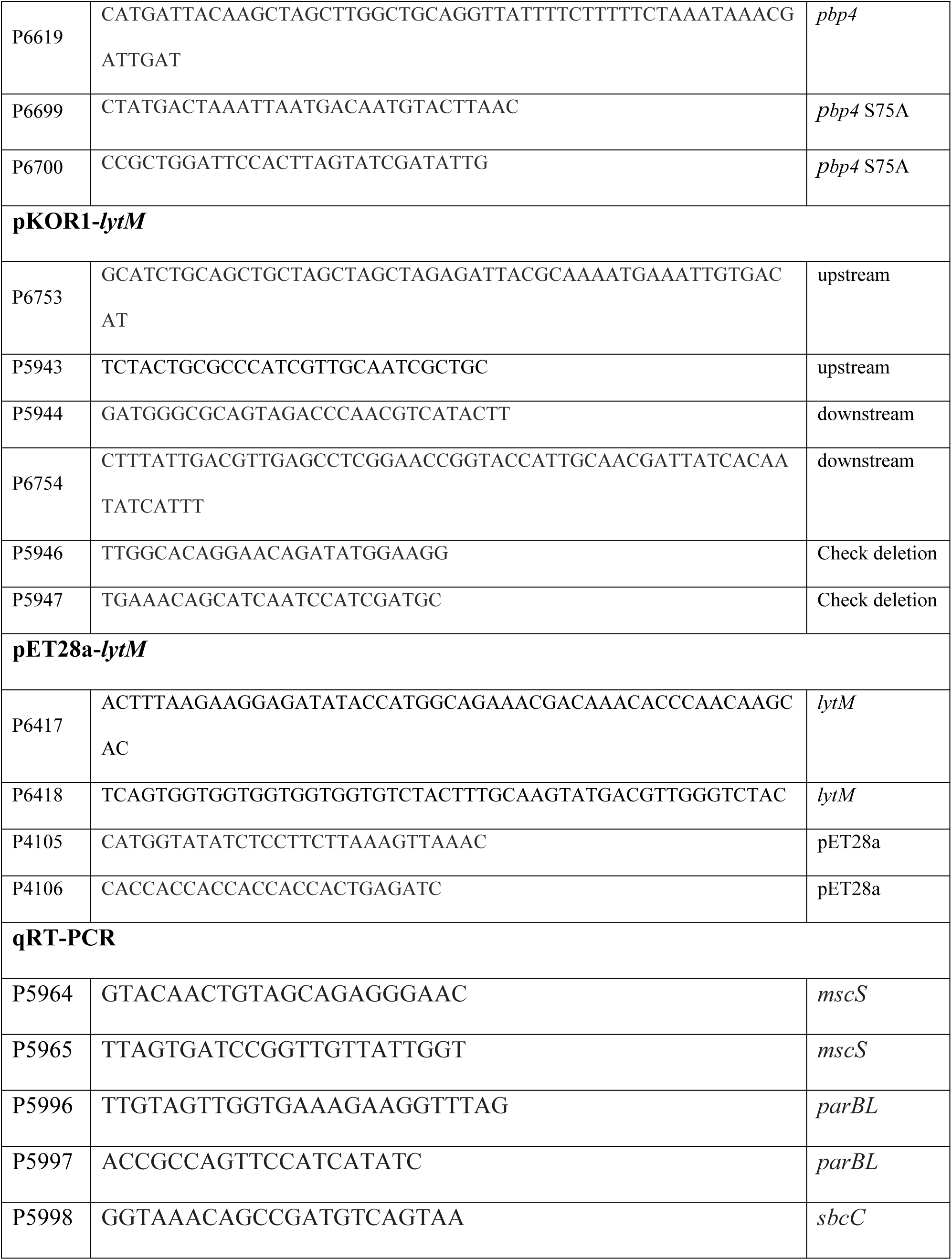
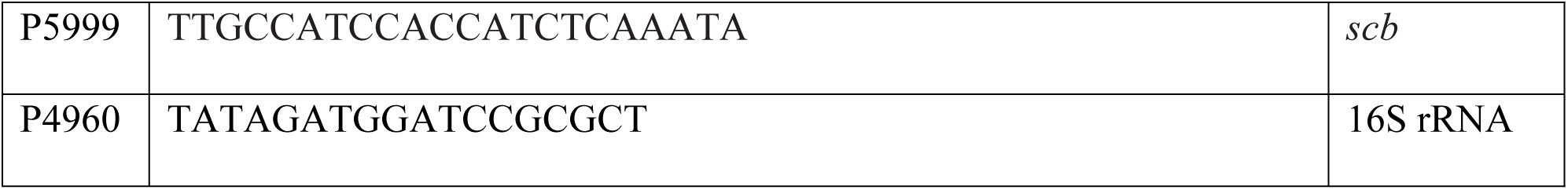
Oligonucleotides used in this study.

### Plasmid construction for ATc-inducible transcription

The plasmids pOS1-Pitet-*parBL-mscS* and pOS1-Pitet-*parBL* were constructed by Gibson assembly. The pOS1 vector was digested with *EcoR*I and *BamH*I. The anhydrotetracycline-inducible promoter Pitet and the *tetR* repressor were PCR-amplified from pYJ335 using primers P6331/P6332 (Table 3). The *parBL-mscS* and *parBL* fragments were amplified from USA300 genomic DNA using primer pairs P6333/P6334 and P6333/P6414 (Table 3), respectively, with Phusion DNA polymerase (NEB). The digested vector and PCR products were purified using a QIAquick PCR Purification Kit (Qiagen) and assembled by Gibson assembly.

The plasmid pOS1-Pitet-*mscS* was derived from pOS1-Pitet-*parBL-mscS* by deleting *parBL*. Deletion was performed by PCR amplification using primers P6415/P6416, followed by *Dpn*I treatment, purification, phosphorylation with T4 polynucleotide kinase (NEB), and ligation with T4 DNA ligase (NEB).

The PBP4-expression plasmid pOS1-Pitet-*pbp4* was derived from pOS1-Pitet-*parBL*. The *pbp4* gene, including its ribosome-binding sequence, was amplified using primers P6618/P6619 and Phusion DNA polymerase (NEB). pOS1-Pitet-*parBL* was digested with *EcoR*I and *Sal*I and assembled with the *pbp4* PCR product by Gibson assembly.

To construct pOS1-Pitet-*pbp4* S75A, pOS1-Pitet-*pbp4* was amplified using primers P6699/P6700 and Phusion DNA polymerase (NEB). The PCR product was treated with *Dpn*I (NEB), purified, phosphorylated with T4 polynucleotide kinase (NEB), and ligated with T4 DNA ligase (NEB).

All plasmid constructs were transformed into *E. coli* DH5α and verified by plasmid sequencing (Plasmidsaurus, Inc.). Verified plasmids were electroporated into *S. aureus* RN4220 and subsequently introduced into the target *S. aureus* strains by φ85-mediated transduction.

### Generation of deletion mutants

To delete *ugtP* in USA300, pCasSA-v2-*ugtP* was constructed as previously described (40). Spacer primers P5288/P5289 were inserted into pCasSA-v2 by Golden Gate assembly. The resulting plasmid was digested with *Bgl*I and *Pvu*I. Upstream and downstream homologous regions were PCR-amplified using primer pairs P5290/P5291 and P5292/P5293, respectively. The PCR products were then assembled with the *Bgl*I/*Pvu*I-digested plasmid by Gibson assembly. The reaction product was transformed into *E. coli* DH5α. The verified plasmid was electroporated into *S. aureus* RN4220 and then transduced into USA300 using phage φ85. Deletion of *ugtP* was confirmed by PCR amplification of the *ugtP* region from the chromosomal DNA using primers P5826/P5593.

To delete *lytM*, pKOR1-*lytM* was generated. Upstream and downstream homologous fragments were PCR-amplified using primer pairs P6753/P5943 and P5944/P6754, respectively, with Phusion DNA polymerase (NEB). The resulting PCR products were purified and assembled with pKOR1-mcs by Gibson assembly (41). The reaction products were transformed into *E. coli* DH5α. Introduction of the resulting plasmid into the target strains and deletion of *lytM* were performed as previously described (41, 42).

### Western blotting

Western blot analysis was performed as described previously (13). Except for the anti-LTA antibody (clone 55, HyCult Biotechnology), all primary antibodies were generated in-house (14). Blots were developed using SuperSignal West Pico PLUS Chemiluminescent Substrate (Thermo Fisher Scientific) and imaged with an AZI400 imaging system (Azure Biosystems). Each experiment was independently repeated at least twice with consistent results.

### LytM antibody generation

The *lytM* sequence lacking the predicted signal peptide was PCR-amplified from USA300 chromosomal DNA using primers P6417/P6418 and Phusion DNA polymerase (NEB). The pET28a vector was amplified using primers P4105/P4106. Purified PCR products were assembled by Gibson assembly, transformed into *E. coli* DH5α, and verified by plasmid sequencing (Plasmidsaurus Inc.). The verified plasmid was transformed into *E. coli* BL21(DE3) (NEB). Recombinant LytM expression was induced with 1 mM IPTG, and His-tagged LytM was purified by nickel-affinity chromatography (Qiagen).

Purified LytM protein was administered via i.m. injection to 12 C57BL/6 mice, three males and three females, at 50 μg per injection for three injections at two-week intervals. One week after the final injection, mice were euthanized, blood was collected by cardiac puncture, and serum was prepared for antibody use.

All animal procedures were conducted in accordance with the Guide for the Care and Use of Laboratory Animals of the National Institutes of Health. The animal protocol, NW-48, was approved by the Committee on the Ethics of Animal Experiments of the Indiana University School of Medicine-Northwest. All efforts were made to minimize animal suffering.

### Determination of minimum inhibitory concentration

Minimum inhibitory concentrations (MICs) were determined using the broth microdilution method according to Clinical and Laboratory Standards Institute (CLSI) guidelines (43). When applicable, MICs were also assessed using MIC test strips (Liofilchem). For strip assays, cultures were grown in TSB to an optical density at 600 nm (OD₆₀₀) of 0.5, and 100 μL of culture was spread onto tryptic soy agar (TSA) plates. MIC strips were applied to the agar surface, and plates were incubated at 37°C for 16–18 h. The MIC was recorded as the antibiotic concentration at which the inhibition ellipse intersected the strip.

### RNA-seq analysis

Test strains were grown in TSB to an OD₆₀₀ of 1.0 and harvested by centrifugation. Cell pellets were lysed in TRIzol reagent (Invitrogen), and total RNA was extracted according to the manufacturer’s instructions. Purified RNA samples were submitted to the Center for Medical Genomics at Indiana University School of Medicine for sequencing. The RNA-seq data were deposited in the Gene Expression Omnibus (GEO) database under accession number GSE330792.

### Quantitative reverse transcription PCR

Test strains were grown in TSB to an OD₆₀₀ of 1.0. Cells were harvested by centrifugation and lysed with lysostaphin (50 μg/mL) for 15 min at 37°C. Total RNA was purified using the RNeasy Mini Kit (Qiagen, catalog no. 74134) according to the manufacturer’s instructions. Complementary DNA (cDNA) was synthesized from purified RNA using random hexamer primers and the High-Capacity cDNA Reverse Transcription Kit (Applied Biosystems). Quantitative PCR was performed using SYBR Green PCR Master Mix (Applied Biosystems) and gene-specific primers listed in Table 3. Each experiment was performed in triplicate, and 16S rRNA was used as the internal reference gene. Relative transcript levels were calculated using the comparative C□ method (2⁻ΔΔ□) (44).

### Measurement of intracellular potassium

Test strains were grown in TSB to an OD₆₀₀ of 1.0. Cells from 1 mL of culture were harvested by centrifugation, and the supernatant was removed. Cell pellets were resuspended in 1 mL of deionized water and treated with lysostaphin (50 μg/mL) for 15 min at 37°C. Cell lysates were then sonicated for three cycles of 15 s, with 15-s intervals, using a Vibra-Cell sonicator (Sonics and Materials, Inc.). Potassium concentrations were determined using a flame photometer, model FF-200 (Cole-Parmer), and quantified by comparison with a standard curve generated from KCl solutions.

### Measurement of intracellular c-di-AMP levels

Intracellular c-di-AMP levels were measured using either a competitive enzyme-linked immunosorbent assay (ELISA) kit (Cayman Chemical) (45, 46) or a *kimA* riboswitch reporter plasmid (35). For ELISA-based measurements, test strains were grown in TSB to an OD₆₀₀ of 1.0. Cells were harvested by centrifugation, treated with lysostaphin (50 μg/mL), and then sonicated. Lysates were centrifuged at 13,000 rpm for 20 min at 4°C, and the resulting supernatants were used for c-di-AMP quantification according to the manufacturer’s instructions.

For the riboswitch reporter assay, the reporter plasmid pCN38-*kimA-yfp* and the control plasmid pCN38-*yfp* were introduced into the WT strain, LTA synthesis mutants, and *gdpP* suppressor mutants. Overnight cultures of the test strains were diluted 1:100 into 150 μL of TSB supplemented with chloramphenicol (10 μg/mL) in black-walled 96-well plates. Plates were incubated in an EnSpire plate reader (PerkinElmer) at 37°C for 18 h with shaking. OD₆₀₀ and fluorescence were measured, with fluorescence detected at an excitation wavelength of 488 nm and an emission wavelength of 535 nm. Fluorescence values were normalized to OD₆₀₀.

Because binding of c-di-AMP to the *kimA* riboswitch represses *yfp* transcription, intracellular c-di-AMP levels are inversely correlated with fluorescence intensity. To express the reporter output as a value that positively correlates with intracellular c-di-AMP abundance, relative c-di-AMP levels were calculated as follows:

Arbitrary unit (AU) = normalized fluorescence from pCN38-*yfp* / normalized fluorescence from pCN38-*kimA-yfp*

All measurements were performed in triplicate.

### β-Galactosidase assay

β-Galactosidase assays were performed as described previously (47), with minor modifications. Overnight cultures were diluted 1:100 into TSB and grown at 37°C with shaking at 200 rpm to the exponential phase. LacZ activity was normalized to OD₆₀₀. Each assay was performed independently at least twice, with consistent results.

### Statistical analysis

Statistical analyses were performed using GraphPad Prism version 10 (GraphPad Software, San Diego, CA, USA). The specific statistical tests used are indicated in the corresponding figure legends.

## Acknowledgments

None

## Supporting Information

**S1 Table. Mutations in the suppressor mutants.**

**S1 Fig. The positions of mutated amino acids in RNA polymerase.** The positions of amino acid substitutions identified in RNA polymerase suppressor mutants are mapped onto the *E. coli* RNA polymerase structure (PDB ID: 6CUX), which was used as a structural model because of the high conservation between *E. coli* and *S. aureus* RNA polymerases. The corresponding residues are indicated by green circles beneath each position marker. The mapped *E. coli* residues corresponding to the mutated *S. aureus* residues are RpoB P519, RpoC G729, RpoC Y1232, and RpoD V579. The structure was visualized using Molstar (molstar.org/viewer).

**S2 Fig. *sbcC* does not contribute to the beta-lactam sensitivity of the *ugtP* mutant.** *S. aureus* USA300 wild-type (WT), the *ugtP* deletion mutant (Δ*ugtP*), and the *ugtP sbcC* double mutant (Δ*ugtP:sbcC*) were spread on TSA plates. Oxacillin MIC test strips were applied, and the plates were incubated overnight at 37°C. The minimum inhibitory concentration (MIC) is indicated below each plate image.

**S3 Fig. Reduced activity of the *parBL* promoter increases oxacillin resistance of USA300Δ*ugtP*.** The native *parBL* promoter was replaced with the IPTG-inducible *Pspac* promoter using the pMutin-HA plasmid in *S. aureus* USA300 wild-type (WT) and Δ*ugtP* strains. The resulting strains were spread on TSA plates, oxacillin MIC test strips were applied, and the plates were incubated overnight at 37°C. The minimum inhibitory concentration (MIC) is indicated below each plate image.

**S4 Fig. Overexpression of individual *parBL* or *mscS* genes does not substantially affect oxacillin resistance in suppressor mutants.** Transcription of either *parBL* or *mscS* was induced from the indicated multicopy plasmid by the addition of anhydrotetracycline at 100 ng/mL. Oxacillin MICs were then determined in liquid culture. – ATc, no anhydrotetracycline; + ATc, with anhydrotetracycline.

**S5 Fig. Co-overexpression of *parBL* and *mscS* sensitizes RNA polymerase suppressor mutants to oxacillin.** The indicated suppressor mutants carrying pOS1-Pitet-*parBL-mscS* were grown to the exponential phase and spread on TSA plates. Oxacillin MIC test strips were applied, and the plates were incubated overnight at 37°C. The oxacillin MIC is indicated below each plate image.

**S6 Fig. PBP4 expression from pOS1-Pitet-*pbp4*.** The indicated strains were grown in TSB to the exponential phase. Anhydrotetracycline was then added to a final concentration of 100 ng/mL, and cultures were incubated for an additional 3 h at 37°C. Cells were collected, normalized by OD₆₀₀, and analyzed by Western blotting using an anti-PBP4 antibody. Sortase A (SrtA) was detected with an anti-SrtA antibody and used as a loading control. Relative PBP4 abundance was calculated by normalizing the PBP4 signal to the corresponding SrtA signal using ImageJ.

**S7 Fig. PBP4 overexpression inhibits the growth of *S. aureus*. (A) Growth inhibition of wild-type strains in TSB.** MW2 and USA300 wild-type strains carrying pOS1-Pitet-*pbp4* were grown at 37°C, and growth was monitored using a Bioscreen C system. (B) Growth inhibition of wild-type strains and LTA synthesis mutants on TSA plates. Strains were grown in TSB to the exponential phase, serially diluted from 10⁰ to 10⁻⁵, and spotted onto TSA plates containing chloramphenicol at 10 μg/mL in the absence (–ATc) or presence (+ATc) of anhydrotetracycline at 100 ng/mL. Plates were incubated overnight at 37°C.

**S8 Fig. *parBL-mscS* transcription does not affect *pbp4* transcription.** The indicated strains were grown to the exponential phase, and *pbp4* transcript levels were quantified by qRT-PCR. Transcript abundance was normalized to 16S rRNA, and values are expressed relative to the USA300 wild-type strain, which was set to 100%. Data represent the mean ± standard deviation from three independent biological replicates.

**S9 Fig. LytM has a minor effect on oxacillin resistance in wild-type strains and LTA synthesis mutants.** The effect of *lytM* deletion on oxacillin resistance was assessed by measuring MICs in liquid culture (A) and on TSA plates using oxacillin MIC test strips (B). The yellow numbers shown on the plate images indicate the oxacillin MICs. –, no *lytM* deletion; Δ*lytM*, *lytM* deletion.

**S10 Fig. ParB and ParBL are structurally distinct.** (A) AlphaFold-predicted structures of ParB and ParBL. (B) CLUSTAL multiple sequence alignment of ParB and ParBL. The structural models and sequence alignment indicate that ParBL is distinct from canonical ParB, despite its annotation as a ParB-like protein.

## Data Availability

RNA-seq data have been deposited in GEO under accession number GSE330792. All other relevant data are within the manuscript and its Supporting Information files.

## Author Contributions

Conceptualization: TB.

Formal analysis: MS, AK, TB.

Funding acquisition: TB.

Investigation: MS, AK, TB.

Methodology: MS, AK.

Project administration: TB.

Supervision: TB.

Validation: MS, AK.

Visualization: MS, AK.

Writing – original draft: TB.

Writing – review & editing: MS, AK, TB.

## References

1. Gordon RJ, Lowy FD. Pathogenesis of methicillin-resistant *Staphylococcus aureus* infection. Clin Infect Dis. 2008;46 Suppl 5:S350–9.

2. Fuda C, Suvorov M, Vakulenko SB, Mobashery S. The basis for resistance to beta-lactam antibiotics by penicillin-binding protein 2a of methicillin-resistant *Staphylococcus aureus*. J Biol Chem. 2004;279(39):40802–6.

3. Utsui Y, Yokota T. Role of an altered penicillin-binding protein in methicillin- and cephem-resistant *Staphylococcus aureus*. Antimicrob Agents Chemother. 1985;28(3):397–403.

4. Mahasenan KV, Molina R, Bouley R, Batuecas MT, Fisher JF, Hermoso JA, et al. Conformational Dynamics in Penicillin-Binding Protein 2a of Methicillin-Resistant *Staphylococcus aureus*, Allosteric Communication Network and Enablement of Catalysis. J Am Chem Soc. 2017;139(5):2102–10.

5. Dordel J, Kim C, Chung M, Pardos de la Gandara M, Holden MT, Parkhill J, et al. Novel determinants of antibiotic resistance: identification of mutated loci in highly methicillin-resistant subpopulations of methicillin-resistant *Staphylococcus aureus*. MBio. 2014;5(2):e01000.

6. Schneewind O, Missiakas D. Lipoteichoic Acid Synthesis and Function in Gram-Positive Bacteria. In: Geiger O, editor. Biogenesis of Fatty Acids, Lipids and Membranes. Cham: Springer International Publishing; 2016. p. 1-18.

7. Grundling A, Schneewind O. Synthesis of glycerol phosphate lipoteichoic acid in *Staphylococcus aureus*. PNAS. 2007;104(20):8478–83.

8. Percy MG, Grundling A. Lipoteichoic acid synthesis and function in gram-positive bacteria. Annu Rev Microbiol. 2014;68:81–100.

9. Kiriukhin MY, Debabov DV, Shinabarger DL, Neuhaus FC. Biosynthesis of the glycolipid anchor in lipoteichoic acid of *Staphylococcus aureus* RN4220: role of YpfP, the diglucosyldiacylglycerol synthase. J Bacteriol. 2001;183(11):3506–14.

10. Grundling A, Schneewind O. Genes required for glycolipid synthesis and lipoteichoic acid anchoring in *Staphylococcus aureus*. J Bacteriol. 2007;189(6):2521–30.

11. Koch HU, Haas R, Fischer W. The role of lipoteichoic acid biosynthesis in membrane lipid metabolism of growing *Staphylococcus aureus*. Eur J Biochem. 1984;138(2):357–63.

12. Hesser AR, Matano LM, Vickery CR, Wood BM, Santiago AG, Morris HG, et al. The length of lipoteichoic acid polymers controls *Staphylococcus aureus* cell size and envelope integrity. J Bacteriol. 2020;202(16):e00149–20.

13. Yeo WS, Jeong B, Ullah N, Shah MA, Ali A, Kim KK, et al. Ftsh Sensitizes Methicillin-Resistant *Staphylococcus aureus* to beta-Lactam Antibiotics by Degrading YpfP, a Lipoteichoic Acid Synthesis Enzyme. Antibiotics (Basel). 2021;10(10).

14. Kanampalliwar A, Shah M, Park Y, Jeong B, Lawson PA, Bell M, et al. The role of lipoteichoic acid in *Staphylococcus aureus* cell wall integrity. bioRxiv. 2025. **doi:** 10.1101/2025.01.16.633316

15. Pinho MG, de Lencastre H, Tomasz A. An acquired and a native penicillin-binding protein cooperate in building the cell wall of drug-resistant staphylococci. Proc Natl Acad Sci U S A. 2001;98(19):10886–91.

16. Hamilton SM, Alexander JAN, Choo EJ, Basuino L, da Costa TM, Severin A, et al. High-Level Resistance of *Staphylococcus aureus* to beta-Lactam Antibiotics Mediated by Penicillin-Binding Protein 4 (PBP4). Antimicrob Agents Chemother. 2017;61(6):e02727–16.

17. Basuino L, Jousselin A, Alexander JAN, Strynadka NCJ, Pinho MG, Chambers HF, et al. PBP4 activity and its overexpression are necessary for PBP4-mediated high-level beta-lactam resistance. J Antimicrob Chemother. 2018.

18. Dengler Haunreiter V, Tarnutzer A, Bar J, von Matt M, Hertegonne S, Andreoni F, et al. C-di-AMP levels modulate *Staphylococcus aureus* cell wall thickness, response to oxidative stress, and antibiotic resistance and tolerance. Microbiol Spectr. 2023;11(6):e0278823.

19. Corrigan RM, Abbott JC, Burhenne H, Kaever V, Grundling A. c-di-AMP is a new second messenger in *Staphylococcus aureus* with a role in controlling cell size and envelope stress. PLoS Pathog. 2011;7(9):e1002217.

20. Dengler V, McCallum N, Kiefer P, Christen P, Patrignani A, Vorholt JA, et al. Mutation in the C-di-AMP cyclase dacA affects fitness and resistance of methicillin resistant *Staphylococcus aureus*. PLoS One. 2013;8(8):e73512.

21. Ba X, Kalmar L, Hadjirin NF, Kerschner H, Apfalter P, Morgan FJ, et al. Truncation of GdpP mediates beta-lactam resistance in clinical isolates of *Staphylococcus aureus*. J Antimicrob Chemother. 2019;74(5):1182–91.

22. Poon R, Basuino L, Satishkumar N, Chatterjee A, Mukkayyan N, Buggeln E, et al. Loss of GdpP Function in *Staphylococcus aureus* Leads to beta-Lactam Tolerance and Enhanced Evolution of beta-Lactam Resistance. Antimicrob Agents Chemother. 2022;66(2):e0143121.

23. Pozzi C, Waters EM, Rudkin JK, Schaeffer CR, Lohan AJ, Tong P, et al. Methicillin resistance alters the biofilm phenotype and attenuates virulence in *Staphylococcus aureus* device-associated infections. PLoS Pathog. 2012;8(4):e1002626.

24. Griffiths JM, O’Neill AJ. Loss of function of the gdpP protein leads to joint beta-lactam/glycopeptide tolerance in *Staphylococcus aureus*. Antimicrob Agents Chemother. 2012;56(1):579–81.

25. Kim JH, Lee Y, Kim I, Chang J, Hong S, Lee NK, et al. Reducing Peptidoglycan Crosslinking by Chemical Modulator Reverts beta-lactam Resistance in Methicillin-Resistant *Staphylococcus aureus*. Adv Sci (Weinh). 2024;11(28):e2400858.

26. Stevens CE, Deventer AT, Johnston PR, Lowe PT, Boraston AB, Hobbs JK. *Staphylococcus aureus* COL: An Atypical Model Strain of MRSA That Exhibits Slow Growth and Antibiotic Tolerance due to a Mutation in PRPP Synthetase. Mol Microbiol. 2025;124(3):189–203.

27. Bae T, Banger AK, Wallace A, Glass EM, Aslund F, Schneewind O, et al. *Staphylococcus aureus* virulence genes identified by bursa aurealis mutagenesis and nematode killing. Proc Natl Acad Sci U S A. 2004;101(33):12312–7.

28. Bose JL, Fey PD, Bayles KW. Genetic tools to enhance the study of gene function and regulation in *Staphylococcus aureus*. Appl Environ Microbiol. 2013.

29. Peyronnet R, Tran D, Girault T, Frachisse JM. Mechanosensitive channels: feeling tension in a world under pressure. Front Plant Sci. 2014;5:558.

30. Gundlach J, Herzberg C, Kaever V, Gunka K, Hoffmann T, Weiss M, et al. Control of potassium homeostasis is an essential function of the second messenger cyclic di-AMP in Bacillus subtilis. Sci Signal. 2017;10(475).

31. Zarrella TM, Metzger DW, Bai G. Stress Suppressor Screening Leads to Detection of Regulation of Cyclic di-AMP Homeostasis by a Trk Family Effector Protein in Streptococcus pneumoniae. J Bacteriol. 2018;200(12).

32. Corrigan RM, Campeotto I, Jeganathan T, Roelofs KG, Lee VT, Grundling A. Systematic identification of conserved bacterial c-di-AMP receptor proteins. Proc Natl Acad Sci U S A. 2013;110(22):9084–9.

33. Moscoso JA, Schramke H, Zhang Y, Tosi T, Dehbi A, Jung K, et al. Binding of Cyclic Di-AMP to the *Staphylococcus aureus* Sensor Kinase KdpD Occurs via the Universal Stress Protein Domain and Downregulates the Expression of the Kdp Potassium Transporter. J Bacteriol. 2015;198(1):98–110.

34. Schuster CF, Bellows LE, Tosi T, Campeotto I, Corrigan RM, Freemont P, et al. The second messenger c-di-AMP inhibits the osmolyte uptake system OpuC in *Staphylococcus aureus*. Sci Signal. 2016;9(441):ra81.

35. Kviatkovski I, Zhong Q, Vaidya S, Grundling A. Identification of novel genetic factors that regulate c-di-AMP production in *Staphylococcus aureus* using a riboswitch-based biosensor. mSphere. 2024;9(10):e0032124.

36. Kreiswirth BN, Lofdahl S, Betley MJ, O’Reilly M, Schlievert PM, Bergdoll MS, et al. The toxic shock syndrome exotoxin structural gene is not detectably transmitted by a prophage. Nature. 1983;305(5936):709–12.

37. Jeong DW, Cho H, Lee H, Li C, Garza J, Fried M, et al. Identification of P3 promoter and distinct roles of the two promoters of the SaeRS two-component system in *Staphylococcus aureus*. J Bacteriol. 2011;193(18):4672–84.

38. Bubeck Wardenburg J, Williams WA, Missiakas D. Host defenses against *Staphylococcus aureus* infection require recognition of bacterial lipoproteins. Proc Natl Acad Sci U S A. 2006;103(37):13831–6.

39. Gibson DG, Young L, Chuang RY, Venter JC, Hutchison CA, 3rd, Smith HO. Enzymatic assembly of DNA molecules up to several hundred kilobases. Nat Methods. 2009;6(5):343–5.

40. Chen W, Zhang Y, Yeo WS, Bae T, Ji Q. Rapid and Efficient Genome Editing in *Staphylococcus aureus* by Using an Engineered CRISPR/Cas9 System. J Am Chem Soc. 2017;139(10):3790–5.

41. Jeong B, Shah MA, Roh E, Kim K, Park I, Bae T. *Staphylococcus aureus* Does Not Synthesize Arginine from Proline under Physiological Conditions. J Bacteriol. 2022;204(6):e0001822.

42. Bae T, Schneewind O. Allelic replacement in *Staphylococcus aureus* with inducible counter-selection. Plasmid. 2006;55(1):58–63.

43. CLSI. Performance standards for antimicrobial susceptibility testing. CLSI approved standard M100-S15: Clinical and Laboratory Standards Institute, Wayne, PA; 2005.

44. Schmittgen TD, Livak KJ. Analyzing real-time PCR data by the comparative C(T) method. Nat Protoc. 2008;3(6):1101–8.

45. Underwood AJ, Zhang Y, Metzger DW, Bai G. Detection of cyclic di-AMP using a competitive ELISA with a unique pneumococcal cyclic di-AMP binding protein. J Microbiol Methods. 2014;107:58–62.

46. Bowman L, Zeden MS, Schuster CF, Kaever V, Grundling A. New Insights into the Cyclic Di-adenosine Monophosphate (c-di-AMP) Degradation Pathway and the Requirement of the Cyclic Dinucleotide for Acid Stress Resistance in *Staphylococcus aureus*. J Biol Chem. 2016;291(53):26970–86.

47. Sun F, Zhou L, Zhao BC, Deng X, Cho H, Yi C, et al. Targeting MgrA-mediated virulence regulation in *Staphylococcus aureus*. Chem Biol. 2011;18(8):1032–41.

